# Public perception of dog emotional and motivational states in videos: A cross-sectional analysis

**DOI:** 10.1101/2025.02.28.640863

**Authors:** Lauren E. Samet, Jana M. Muschinski, Naomi D. Harvey, Kassandra Giragosian, Melissa Upjohn, Jane Murray, Sara Owczarczak-Garstecka

## Abstract

Recognition and interpretation of dogs’ emotional and motivational states from visual behavioural signs are important for public safety and dog welfare. This study used an online survey to explore the ability of members of the public (n = 4,133) to recognise the underlying emotional or motivational states of dogs in silent videos (n=30). Participants scored each video for nine pre-determined emotional and motivational states on a scale from 0 to 15 and rated the relative difficulty of scoring each video. Participants could also select “I am uncertain” for individual states which translated to missing values. Public scores were compared with those of eleven dog behaviour experts. The states “nervous/anxious”, “stressed”, “relaxed”, “comfortable”, “playful”, “interested/curious”, “excited”, and “frustrated” showed high inter-expert agreement and were used in further analysis. “Boredom” was removed due to low inter-expert agreement. Principal components and cluster analyses on both datasets were used to collapse categories into two dimensions, identify groupings and compare overall perception. Results indicate similarity in perception of underlying states between public and experts. Correlation between expert difficulty rating, and both inter-expert agreement and public accuracy, indicates that experts effectively assessed the relative difficulty of determining underlying state. Members of the public perceived playful, excited, and curious dogs as easier to interpret than anxious and stressed dogs; however, this was not reflected in how accurately they scored videos (i.e., how different a participant’s scores were from the expert scores) and instead was reflected by how likely a participant was to score a video in full, rather than selecting that they were “uncertain” in response to any of the listed states. Findings of this study inform human behaviour change interventions to improve public interpretation of dog emotional and motivational states.

## 1 Introduction

Accurate identification and interpretation of dog behaviour is vital for positive human-dog interactions, public safety, and dog welfare. Poor recognition of signs associated with distress and anxiety in dogs can lead to inappropriate responses and the escalation of behaviours. This can lead to behaviours that pose a problem for the dog’s guardian or the public, or may increase the risk of relinquishment or euthanasia (Boyd et al., 2018; Hargrave, 2015; Kwan & Bain, 2013; Overall, 2013). Additionally, enhancing human understanding of dog behaviours that signpost unmet welfare needs can prompt effective remedial action (Pegram et al., 2021). Thus, it is crucial that the public can accurately identify and interpret dog behaviour to mitigate negative consequences for both dogs and humans.

Emotions are multicomponent (physiological, behavioural, and cognitive) subjective responses to a stimulus or event that have salience to the individual. Emotions are always valenced (have pleasant or unpleasant hedonic tone) and can vary in activation/arousal and duration/persistence (Paul & Mendl, 2018). Motivational states are driven by internal physiological changes that energise an organism to achieve its desired goal, e.g., quenching hunger, thirst or caring for offspring (Alarcón, 2020). Emotions and motivations are often discussed together as emotional and motivational “states” due to their synergistic links between physiology and neural pathways. Additionally, pursuing motivating behaviour can feed into emotional responses – behaviours can be driven by stimulus-elicited emotions as well as goal-achieving motivations (Berridge, 2018).

Behaviours, or groups of behaviours, can correspond to specific emotional and motivational states (e.g., Tami & Gallagher, 2009; Walker et al., 2010). Some dog behaviours, such as lowered postures (including body, tail, ears) (Beerda et al., 1997), excessive salivation, vocalisations, hiding, pacing, panting, remaining near the owner, trembling (Dreschel & Granger, 2005), freezing, retreating, flinching and paw lifting (Flint et al., 2018) can indicate distress (negative/detrimental experience). However, other behaviours are only equivocally associated with emotional or motivational states (e.g., paw raising may be associated with anxiety or excitement) and some with ambivalence too (e.g., panting). This poses a challenge for lay people when interpreting the meaning of different dog behaviours (Pastore et al., 2011).

Members of the public perceive dogs differently depending on the behaviour, emotional, or motivational state they are asked to identify, and show greater accuracy assigning some states over others (e.g., Bloom & Friedman, 2013; Lakestani et al., 2014; Törnqvist et al., 2023). Studies found that the public were better at identifying dogs the researchers had labelled as “aggressive”/“angry” from photos, videos, and audio, compared to dogs displaying states such as “fearful”, “sad”, or “surprised” (Amici et al., 2019; Bloom & Friedman, 2013; Lakestani et al., 2014). This pattern was consistent for both those experienced and inexperienced with dogs (Bloom & Friedman, 2013), though experience was in some cases found to improve overall performance (Amici et al., 2019). Törnqvist et al. found that adults with dog ownership experience were more likely to classify photos of “aggressive dogs” correctly than of “happy dogs”, while both those with and without ownership experience classified photos of “neutral dogs” more accurately than those of “happy dogs” (2023).

Studies have also reported that “happiness”/“friendliness” and “play” are easier to identify than states such as “fear”, “anxiety”, or “sadness” (Amici et al., 2019; Bloom & Friedman, 2013; Lakestani et al., 2014; Wan et al., 2012), but across studies these results were inconsistent. Tami and Gallagher found that the public had difficulty differentiating between instances of "aggression" and "play" when viewing silent videos (2009). They found instances of “fear”, “friendliness”, “indifference”, and “play solicitation” (c.f. “actual play”) to be easier to interpret (Tami & Gallagher, 2009). This could indicate that vocalizations or combination of visual and vocal cues may be important in the recognition of emotional state. This is further supported by research showing that both adults and children rely on sound cues to identify aggressive behaviour from videos (Lakestani et al., 2014).

The public have also been found to struggle identifying mild or moderate emotional responses (e.g., fear), possibly due to overlooking subtle behavioural signs (Flint et al., 2018; Mariti et al., 2012). Patterns of recognition of emotional and motivational states are nuanced and can lead to conflicting results. For example, some studies reported high accuracy of members of the public’s ability to identify "fear" and "nervousness" (Bahlig-Pieren & Turner, 1999; Diesel et al., 2008; Tami & Gallagher, 2009), while others reported low accuracy (Amici et al., 2019; Bloom & Friedman, 2013; Demirbas et al., 2016; Lakestani et al., 2014). These discrepancies in findings can be partially explained by variation in experimental methodology. Noteworthy distinctions include studies employing videos with and without sound, as well as use of photos of facial expression versus full-body videos. Discrepancies between findings may also be due to small sample sizes (<10 video clips) and assigning one emotional or motivational state to each dog (e.g., Tami & Gallagher, 2009). This study design forces respondents to select a single state, despite behavioural expressivity often conveying more than one emotion, even if one predominates (Panksepp, 2010). Human perceptions of dog behaviours can also be modulated by familiarity with dogs and their behaviour, past interactions, attitudes towards dogs, dog characteristics, and the surrounding environment (Arhant et al., 2010; Jalongo, 2018; Meints & de Keuster, 2009; Owczarczak-Garstecka et al., 2019; Westgarth & Watkins, 2015). This means that differences in sample populations may influence outcomes.

Further inconsistent findings may be partly explained by the wide-ranging terminology applied to emotional and motivational states. Whilst some studies use adjectives, others have applied basic emotions (Ortony & Turner, 1990), valence and arousal measures, or terms relating to qualitative behavioural assessments (QBA) (Wemelsfelder et al., 2001). The public struggle to describe the specific dog behaviours that underpin their interpretation and instead often prefer to refer to dogs’ emotional and motivational states (Demirbas et al., 2016; Oxley et al., 2022; Tami & Gallagher, 2009). Dog trainers also use holistic terminology to describe dog behaviour (Tami & Gallagher, 2009). The use of adjectives (e.g., indifferent, fearful, confident, friendly, playful) as descriptors of behaviours, emotions, and motivations, makes classification of emotions and cross-study comparison challenging. This also underscores the complexities that researchers encounter in defining these concepts.

### Public ability to recognise their own level of knowledge in dog behaviour

Issues relating to the public’s ability to interpret dog behaviour, emotional and motivational states are exacerbated when individuals perceive their skills differently from their actual ability. Kerswell et al. (2009, 2013) found that the public generally reported higher levels of confidence identifying positively valenced emotional and motivational states than negative ones. However, no correlation was found between the self-confidence ratings for different emotional and motivational states and the number of behavioural indicators participants could list for those states, suggesting confidence was potentially unfounded (Kerswell et al., 2009).

Kerswell et al. found dog ownership was correlated with confidence levels, with increased experience potentially leading to overconfidence, resulting in a compromised interpretation of dog body language (2013). Moss and Wright (1987) reported that dog owners indicated they were willing to approach a dog displaying potential warning signs to a closer distance compared to non-dog owners. Similarly, Demirbas et al. (2016) found dog owners were less adept at identifying potentially risky scenarios between children and dogs, than non-dog owners. As evidenced by these studies, confidence may not always correspond with level of understanding. Exploring the relationship between confidence and competence is needed to design effective interventions.

### Study aims

This study assessed the ability of members of the public to interpret underlying emotional and motivational states from videos of dogs using QBA terms, and to evaluate their perceived difficulty of interpretation. Participants were presented with a series of silent videos depicting dog behaviour, and their responses were compared to those of dog behaviour experts. The study aimed to:

1. Compare interpretation of dog emotional and motivational states between members of the public and experts;
2. Identify how challenging different emotional and motivational states in the videos are for both the public and experts to assess;
3. Investigate the accuracy of the public and experts in evaluating the difficulty of interpreting a specific dog’s behaviour and their overall competency.

## 2 Methods

An online survey was used to investigate United Kingdom (UK) and Republic of Ireland (ROI) public perceptions of dog behaviour via the use of thirty silent video clips of dogs, each <30 seconds in length, in a range of common environments displaying a variety of behaviours.

### 2.1 Public survey tool

Data were collected via a self-administered online survey (Table S1 in Supplementary Materials) developed using SmartSurvey™ (https://www.smartsurvey.co.uk/), estimated to take approximately 25 minutes to complete. The survey was tested by Dogs Trust staff members and refined prior to public release. The survey (open from 5 May – 1 July 2021) was promoted via social media platforms, sponsored Facebook advertisements, Dogs Trust e-newsletter, local press coverage across several UK regions, national online press, and emails to consenting participants of previous Dogs Trust research studies. Individuals aged 18 years or older residing in the UK or ROI, irrespective of their experience with dogs or dog ownership status, were eligible to participate. Informed written consent was obtained from all subjects involved in the study using the survey platform.

#### 2.1.1 Dog behaviour videos and randomization

Silent video clips presented to participants were opportunistic recordings of spontaneous dog behaviour within a range of contexts. Most recordings included a single dog, but where multiple dogs (or other animals) were included, the focal dog was specified by the dog’s coat colour or other features. The contexts included dogs interacting with people, animals, other dogs, or environmental stimuli in homes, gardens, and out on walks (off lead and on lead). Each participant was shown five of the 30 videos. Due to software limitations preventing randomization of video assignment and order, six video groups (labelled A – F) of five videos each were created in advance, with each group containing a variety of contexts and behaviours. Each group had a “forward” and “reverse” version, resulting in a total of 12 groups, to allow testing for effects of video order. The 12 video groups were named after the months of the year, and participants were prompted to select the birth month of a friend, family member, or pet, to facilitate random selection of a video group. See Table S2 in Supplementary Materials for a list of video descriptions and groupings.

#### 2.1.2 Video-related questions

To facilitate lay understanding, the term “emotions”, rather than “emotional and motivational states” was used. The emotional and motivational states were chosen based on previously validated QBA terms identified by Arena et al. (2019) and consultations with two dog behaviour experts who viewed the videos. QBA builds on the use of non-specific, anthropomorphic language, using common parlance descriptive terms that include emotional states (such as “relaxed” or “anxious”) to describe animal behaviour and capture an animal’s overall demeanour (Arena et al., 2019; Wemelsfelder et al., 2001). As QBA terms do not require translation into lay terms when accompanied by a suitable definition, this approach can be useful when assessing public understanding of the relationship between observed behaviours and underlying emotional and motivational states, and for comparisons between different groups. “Frustration” was added to the QBA terms because it was a descriptor used repeatedly by the behaviour experts. The final list of included terms was: “nervous/anxious”, “stressed”, “relaxed”, "comfortable”, “frustrated”, “excited”, “playful”, “interested/curious”, and “bored”.

Participants were provided with definitions of each term and were asked to score on a scale from 0 to 15 how strongly they thought the dog in a video was experiencing each one (i.e., “Please rate to what extent you think the dog in the video is experiencing each of the emotional descriptors below”; Table 1). Participants could also select “I am uncertain” instead of providing a score for a given QBA term. After completing the scoring, participants were asked to rate the perceived difficulty of scoring the video (i.e., “Using the scale below, rate how difficult or easy you found it to tell how the dog was feeling in this video” with answers on a five-point Likert-scale from “very difficult” to “very easy”).

**Table 1:**
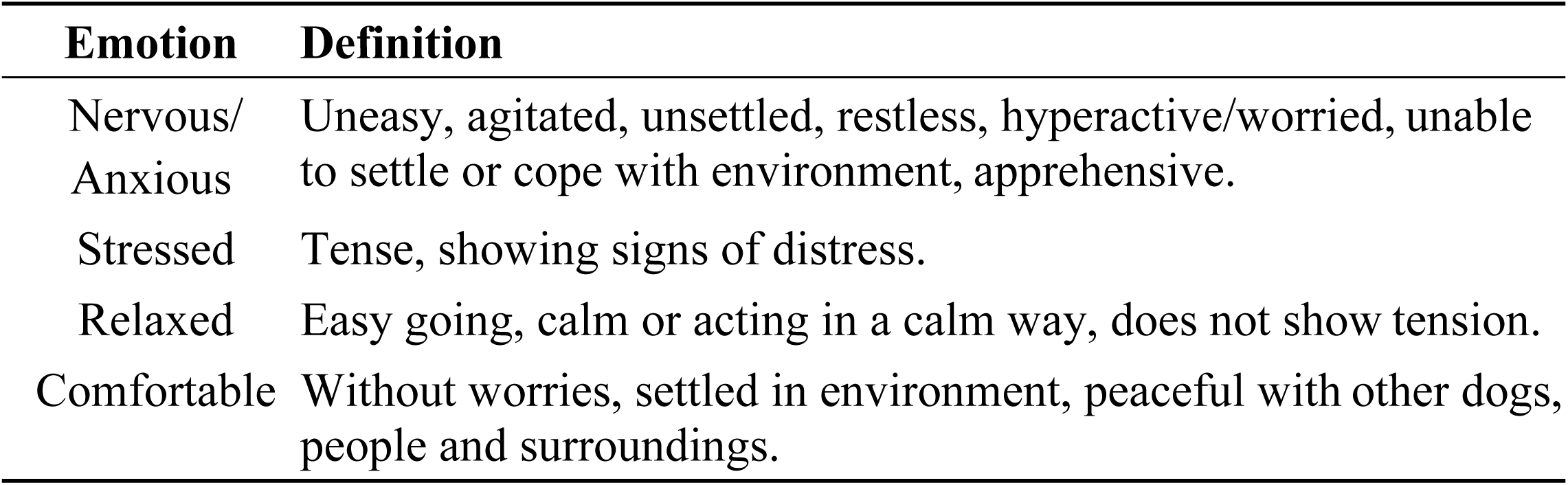
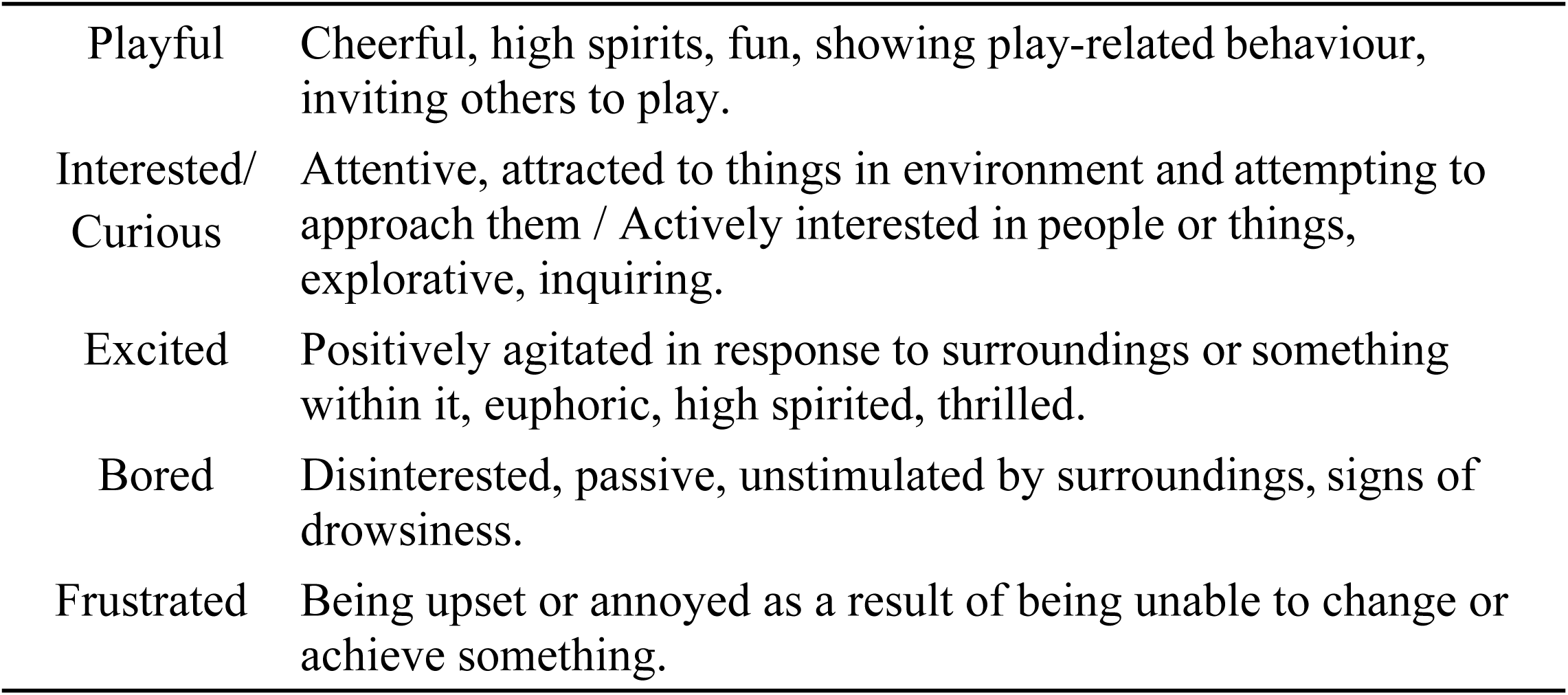
Nine emotional and motivational states (QBA terms) used for scoring the focal dogs within the videos provided in the survey (based on definitions from Arena et al., 2019). These were referred to as “Emotions” to assist understanding.

| Emotion | Definition |
| --- | --- |
| Nervous/<br>Anxious | Uneasy, agitated, unsettled, restless, hyperactive/worried, unable to settle or cope with environment, apprehensive. |
| Stressed | Tense, showing signs of distress. |
| Relaxed | Easy going, calm or acting in a calm way, does not show tension. |
| Comfortable | Without worries, settled in environment, peaceful with other dogs, people and surroundings. |
| Playful | Cheerful, high spirits, fun, showing play-related behaviour, inviting others to play. |
| Interested/<br>Curious | Attentive, attracted to things in environment and attempting to approach them / Actively interested in people or things, explorative, inquiring. |
| Excited | Positively agitated in response to surroundings or something within it, euphoric, high spirited, thrilled. |
| Bored | Disinterested, passive, unstimulated by surroundings, signs of drowsiness. |
| Frustrated | Being upset or annoyed as a result of being unable to change or achieve something. |

#### 2.1.3 Person-specific questions

Participants were asked optional questions about themselves regarding their experience with dogs, education relating to dog training and behaviour, attitudes towards dogs, training beliefs, and demographics (Table S1). Questions related to attitudes towards dogs included a self-assessment of participants’ ability to interpret dog behaviour and body language (i.e., “I feel confident interpreting any dog’s behaviour”, rated on a five-point Likert-scale from “strongly disagree” to “strongly agree”). Several person-specific questions, marked in Table S1, were based on those used in Generation Pup surveys to enable comparisons (Murray et al., 2021). Demographic questions included participants’ gender identity, age range, household income range, and the first part of their postcode (i.e., area and district).

### 2.2 Expert survey tool

Expert survey data (ESD) were collected to establish a baseline for “ideal” question answers for the videos, and to facilitate a comparison with responses from the public. Eleven Dogs Trust staff members with expertise and/or qualifications in dog behaviour answered the same questions for all 30 videos using a separate online survey (also developed within SmartSurvey™). The expert-completed survey replicated the questions posed to the public but omitted the person-specific questions.

### 2.3 Ethics

The study was approved by the Dogs Trust Ethical Review Board (ERB041).

### 2.4 Data cleaning

Public survey data (PSD) were downloaded and identifying participant information removed (e.g., email address). Ahead of the analysis, participants who rated fewer than four out of the five videos in their assigned group were removed to ensure a dataset with minimal missing data. Duplicate responses were identified and removed based on IP address.

### 2.5 Data analysis – an overview

All analyses were conducted using the statistical programming software R version 4.3.2 (2023-10-31 ucrt) (R Core Team, 2023). First, the inter-observer reliability between expert participants was evaluated (2.5.1. Inter-observer reality checks). This process aimed to identify and exclude QBA terms that could not be reliably assessed by experts from the videos, and to discard videos where expert consensus was not reached.

Next, principal component analysis (PCA) and k-means clustering were used to collapse remaining QBA terms into two principal components, and identify videos where experts perceived dogs to exhibit similar emotional or motivational states (2.5.2. PCA and Clustering: Reducing dimensions and identifying video groups). This process was repeated for the public dataset to evaluate the disparity in overall perception of the videos between the public and experts. Finally, the accuracy and perceived difficulty of the public’s scoring were compared across the video clusters. Scoring difficulty was represented by the subjective “difficulty” rating assigned by each participant to each video, whilst scoring accuracy was determined using a calculated difference variable reflecting how an individual scored in relation to the experts (2.5.3. Measuring accuracy and subjective difficulty).

#### 2.5.1 Inter-observer reliability checks

An absolute agreement two-way mixed effects model of Intraclass Correlation Coefficient (ICC; (Koo & Li, 2016)) was completed using the *irr* package (Gamer et al., 2022) to measure the reliability of mean scores per video between the 11 experts. Of the 30 videos, four were removed from further analysis due to a high proportion of missing data, i.e., n ≥ 15 “I am uncertain” of 99 possible scores (Table S2). Three additional videos were removed due to low ICC estimates (≤0.5) indicating low expert agreement. Of the remaining 23 videos, five had ICC estimates >0.75 and 18 had estimates >0.9, indicating high inter-observer agreement.

The same process was completed for each QBA term, for the 23 remaining videos. Eight of nine QBA terms were retained, with ICC estimates >0.9 indicating high inter-observer reliability (Koo & Li, 2016). The QBA term “bored” had an ICC estimate of 0.24 and was removed from further analysis. Videos and QBA terms removed from the expert dataset were also removed from the public dataset.

#### 2.5.2 PCA and clustering: Reducing dimensions and identifying video groups

For the 23 remaining videos, PCA was used to identify overlap in the QBA terms and reduce dimensions. Two separate PCA analyses were carried out: (1) on QBA scores provided by the public, and (2) on expert scores. All PCA analyses were conducted with a correlation matrix of z-score standardized data for each QBA term.

For each analysis, a Kaiser-Meyer-Olkin (KMO) test and Bartlett’s Test of Sphericity were used to confirm sampling adequacy. KMO tests led to the removal of the “frustrated” QBA term for the ESD (Measure of sampling adequacy [MSA] = 0.3 for “frustrated”), but not the PSD (MSA = 0.85 for “frustrated”) (Budaev, 2010). The number of retained components was selected by visual inspection of eigenvalues and scree plots. Eigen value decomposition PCA was completed with two components retained using the *psych* package (Revelle, 2022). Loadings were varimax rotated and rotated component scores calculated for each video. Following PCA, the inter-observer reliability between experts of both rotated components were assessed using a two-way mixed effects absolute agreement multiple raters’ ICC to decide on further steps for video analysis. ICC estimates were greater than 0.9 for both rotated components, indicating high reliability between experts, therefore a by-video mean for each component was taken. The mean rotated component scores (RC1 and RC2) for each video were calculated separately for expert and public scores.

K-means clustering was applied to the results of the expert PCA to identify clusters of videos showing dogs evaluated as exhibiting similar underlying states. K-means clustering is an unsupervised machine learning algorithm which groups points to maximise intra-class similarity while minimising inter-class similarity (MacQueen, 1967). Here, each point represented one of the 23 videos and consisted of the video’s expert mean RC1 and RC2 scores. K was determined in advance using the elbow and silhouette methods (Charrad et al., 2014; Yuan & Yang, 2019). K-means clustering was completed using the *Stats* package (R Core Team and Worldwide Contributors, 2022) with the Hartigan and Wong algorithm (Hartigan & Wong, 1979). Clustering outcomes remained consistent regardless of whether by-video means were computed before or after PCA was performed. The same process was completed for the public PCA results, with each point corresponding to the public mean RC1 and RC2 of one video. This was done to identify any differences in the overall perception of specific videos by experts versus the public (i.e., differences in how videos clustered together).

#### 2.5.3 Measuring accuracy and subjective difficulty

By comparing metrics of difficulty and accuracy across video clusters, it was possible to evaluate which clusters were more likely to pose objective and subjective challenges for the public in scoring, and to determine if participants were adept at evaluating their own ability to interpret dog emotional and motivational states. To achieve this, we measured public accuracy relative to expert scores for each video (Method 1), summarized participants’ certainty during the scoring of each QBA term (Method 2) and described the overall subjective difficulty of a video as perceived by participants (Method 3).

##### Method 1: Objective accuracy – The “difference variable”

The difference variable (DV) was calculated for each public participant’s observations to identify videos where the participant excelled or struggled at scoring, compared to an “ideal” expert score. The “ideal” expert scores were defined as the median of the expert scores for each retained QBA term, for a given video. Medians were used rather than means as they are more resistant to skewing caused by outliers, providing a more robust measure of central tendency. DV scores for each public participants’ observations were defined as the Euclidean distance between the vector of public scores for each of the eight retained QBA terms for a given video, and the vector of the relevant “ideal” expert scores.

##### Method 2: Certainty – “I am uncertain” scores

When identifying the state associated with dogs’ behaviours shown in the videos, public participants and experts were also able to choose the option “I am uncertain” (IAU) rather than scoring a QBA term. An IAU response was used as a proxy measure of difficulty with video interpretation and translated into a missing value when calculating DV.

##### Method 3: Subjective difficulty – Likert difficulty rating

Public participants and experts were asked to rate on a 5-point Likert scale (“very difficult”, “difficult”, “neither easy nor difficult”, “easy”, “very easy”) how difficult they found it to score each video.

#### 2.5.4 Modelling

Modelling was completed using the package *lme4* with maximum likelihood estimation, unless otherwise specified (Bates et al., 2015, 2024). Multiple models/analyses with different outcome variables were constructed (see Table 2).

**Table 2:** Model and analysis comparisons and predictions.

| Research Question | Comparison | Prediction |
| --- | --- | --- |
| **What emotional and motivational states do the public and experts find difficult to assess?** | Video cluster cf. accuracy and subjective difficulty (Models 1 and 3) | Clusters 1 (fear/anxiety) and 2 (play/excitement) were expected to have the lowest DV and difficulty scores, with Cluster 3 (neutral/comfortable) having higher DV scores and difficulty ratings (Amici et al., 2019; Bahlig-Pieren & Turner, 1999; Bloom & Friedman, 2013). |
|  | Rotated components cf. accuracy and subjective difficulty (Models 2, 4, 5, and 6) | It was expected that rotated component (RC) 1 would be positively correlated with DV and difficulty, as positive RC1 scores were associated with more neutral behaviours and should be more difficult to score. It was expected that RC2 would be negatively associated with DV and difficulty, as a high RC2 score was associated with very playful behaviour and members of the public are reported to find identifying “happy” emotions easier compared to fearful, neutral, and sad emotions (Amici et al., 2019).<br>A significant interaction between the two components was expected because a very high RC2 score with a very negative RC1 score would indicate a very mixed, and therefore difficult to interpret, emotional state. |
| **How well do people assess difficulty and their own competence?** | Expert difficulty rating cf. inter-expert agreement estimates (Analysis 7) | It was expected that experts would be good at assessing the difficulty of videos, and therefore a strong negative correlation between experts’ perception of difficulty and the ICC was predicted. |
|  | DV score cf. subjective video public difficulty rating, expert difficulty rating, and self-assigned competence (Models 8, 9, and 10) | A positive correlation was expected between difficulty (both expert and public) and DV score, and potentially a negative correlation between self-assigned competence and DV score. Several studies reported that a higher level of dog experience can be associated with lowered performance in identifying dog emotions (Bloom & Friedman, 2013; Demirbas et al., 2016, but see Amici et al., 2019). |

Models fell into three overall categories, assessing: 1) video difficulty, 2) public scoring accuracy, and 3) the relationship between accuracy, self-assigned competence, and perceived difficulty across video types. In addition to the complete case analyses (Models 1, 2, 3, 4, 8, 10, and analysis 7), a series of missingness models (Models 5, 6, and 9) were run to assess how “I am uncertain” scores impacted analyses. Models 1, 2, 5, 6, and 9 were binary logistic regression models, with Tukey post-hoc test using the *emmeans* package when relevant (Lenth et al., 2018).

Where individuals selected “I am uncertain” rather than a score from 0 to 15 for any of the retained QBA terms, this resulted in a missing value. This led to missing data in models with DV as an outcome. As such, results from Models 3 and 4 are interpreted as referring to accuracy when participants were certain enough to provide a score for all QBA terms, while further models (Models 5, 6, and 9) investigated patterns of missingness. These models were run without the random effect of video ID due to singularity issues when including both video ID and participant ID (Table 3). We did not impute values for unscored QBA terms as these values were not “missing” as in “uncollected” but were the direct result of participants acknowledging that they were uncertain. Instead, to provide a more complete picture of accuracy across clusters and accuracy vs. perceived difficulty, we present both available case analysis and missingness analyses (Models 5, 6, and 9).

**Table 3:** Model and analysis outcomes, fixed effect, and random effects for models described in Table 2.

| Model / Analysis | Outcome | Fixed effects | Random effects |
| --- | --- | --- | --- |
| Model 1 | “Difficult” vs “Easy” | Video cluster (from K-means clustering) | Video ID, Participant ID |
| Model 2 | “Difficult” vs “Easy” | Rotated component (RC) 1, RC2, interaction between RC1 and RC2 | Video ID, Participant ID |
| Model 3 | Log(DV) | Video cluster | Video ID, Participant ID |
| Model 4 | Log(DV) | RC1, RC2, interaction between RC1 and RC2 | Video ID, Participant ID |
| Model 5 | Missing DV | Video cluster | Participant ID |
| Model 6 | Missing DV | RC1, RC2, interaction between RC1 and RC2 | Participant ID |
| Analysis 7 | Video's ICC coefficient | Average expert difficulty rating (Spearman's Rank Order Correlation analysis) | N/A |
| Model 8 | Log(DV) | Participant's video difficulty rating, self-rated confidence, interaction between difficulty and self-rated confidence | Video ID, Participant ID |
| Model 9 | Missing DV | Participant's video difficulty rating, self-rated confidence, interaction between difficulty and self-rated confidence | Participant ID |
| Model 10 | Log(DV) | Video's avg. expert difficulty rating, participant's self-rated confidence, interaction between avg. expert difficulty rating and participant confidence | Video ID, Participant ID |

Ahead of modelling, some variables were transformed. DV was log transformed whenever used as an outcome to ensure normally distributed residuals. When “difficulty” (as rated by the public) was used as a model outcome, it was collapsed into a binary variable (“difficult” and “very difficult” collapsed into “difficult”; “easy” and “very easy” collapsed into “easy”), with “neither difficult nor easy” observations dropped from the analysis. This allowed for the use of binary instead of ordinal logistic regression to avoid violating the assumption of proportional odds when all five difficulty categories were included. Random effects models including video ID, participant ID, and video group and direction (e.g., group B, reversed video order; or group F, forward video order) were used to assess which random effects explained significant variance in the outcome and should be included in further modelling. Video and participant IDs were used as random effects in following mixed effects models, while video group/direction was not as there was no evidence of a group or video order effect (ICC < 0.1).

Using Akaike information criterion (AIC) and Bayesian information criterion (BIC), all models were compared to their associated null model, which contained only the relevant random effects. When models included more than one variable of interest, backwards selection was used, with the final model compared to the null model. Normality of residuals and homoscedasticity were checked visually. Overdispersion was checked using the *blmeco* package (Korner-Nievergelt et al., 2019) and results are reported in Section IV in Supplementary Materials.

## 3 Results

The survey was completed by 5,300 public participants; 4,133 participants answered questions about at least four out of the five videos in their set and were included in analysis. Demographic questions within the survey were non-compulsory (n _total_ refers to total question respondents). Most participants identified as female (n _total_ = 3,868 respondents; n _female_ = 3,354; 86.7%), lived in England (n _total_ = 3,759; n _England_ = 2,932; 78.0%), and were White British (n _total_ = 3,704; n _white-British_ = 3,033; 81.9%). Male respondents were spread evenly across clusters (Cluster 1: 11.4% male, 86.6% female; Cluster 2: 11.5% male, 86.7% female; Cluster 3: 10.7% male, 87.2% female) and across video groups (percentages of male participants between 9.3% and 13.5% for all six video groups).

Of participants who responded to the statement “I feel confident interpreting any dog’s behaviour” (n = 3,805), only 1.9% reported that they completely disagreed and 9.4% reported that they slightly disagreed. A further 11.9% neither agreed nor disagreed, while 55.6% slightly agreed and 21.3% completely agreed. See Supplementary Materials, Section I for additional descriptive statistics on demographic and dog experience questions.

### 3.1 Principal component analysis and clustering

Two components accounted for 81% of variance in the ESD and for 75% of variance in the PSD (Table). Communalities were high for all included QBA terms in the ESD (all communalities >0.7). Communalities were high for all included QBA terms except “frustrated” in the PSD (“frustrated” communality = 0.47, all others >0.7).

**Table 4:** Rotated structure matrix for PCA with varimax rotation for the two-component emotional and motivational state responses for both expert and public data. Values >0.7 in bold.

| QBA Term | Rotated Component 1 |  | Rotated Component 2 |  |
| --- | --- | --- | --- | --- |
|  | Expert | Public | Expert | Public |
| Nervous/Anxious | **-0.742** | **-0.770** | -0.465 | -0.461 |
| Stressed | **-0.711** | **-0.765** | -0.479 | -0.467 |
| Relaxed | **0.920** | **0.795** | 0.130 | 0.306 |
| Comfortable | **0.872** | **0.777** | 0.268 | 0.428 |
| Playful | 0.336 | 0.335 | **0.845** | **0.858** |
| Interested/Curious | 0.170 | 0.216 | **0.842** | **0.828** |
| Excited | 0.343 | 0.106 | **0.886** | **0.926** |
| Frustrated | *Excluded* | -0.674 | *Excluded* | 0.119 |

Scores for both rotated components had ICC estimates greater than 0.9 for inter-expert reliability, therefore means of scores were taken by video to create an RC1 and an RC2 score per video. The same was carried out for the public results; K-means clustering on the by-video means resulted in the same clustering patterns for both the PSD and ESD (Figure 1). The sample means and standard deviations of expert scoring for each QBA term per cluster are shown in Table to demonstrate the key emotional and motivational state characteristics of each cluster.

**Figure 1:**
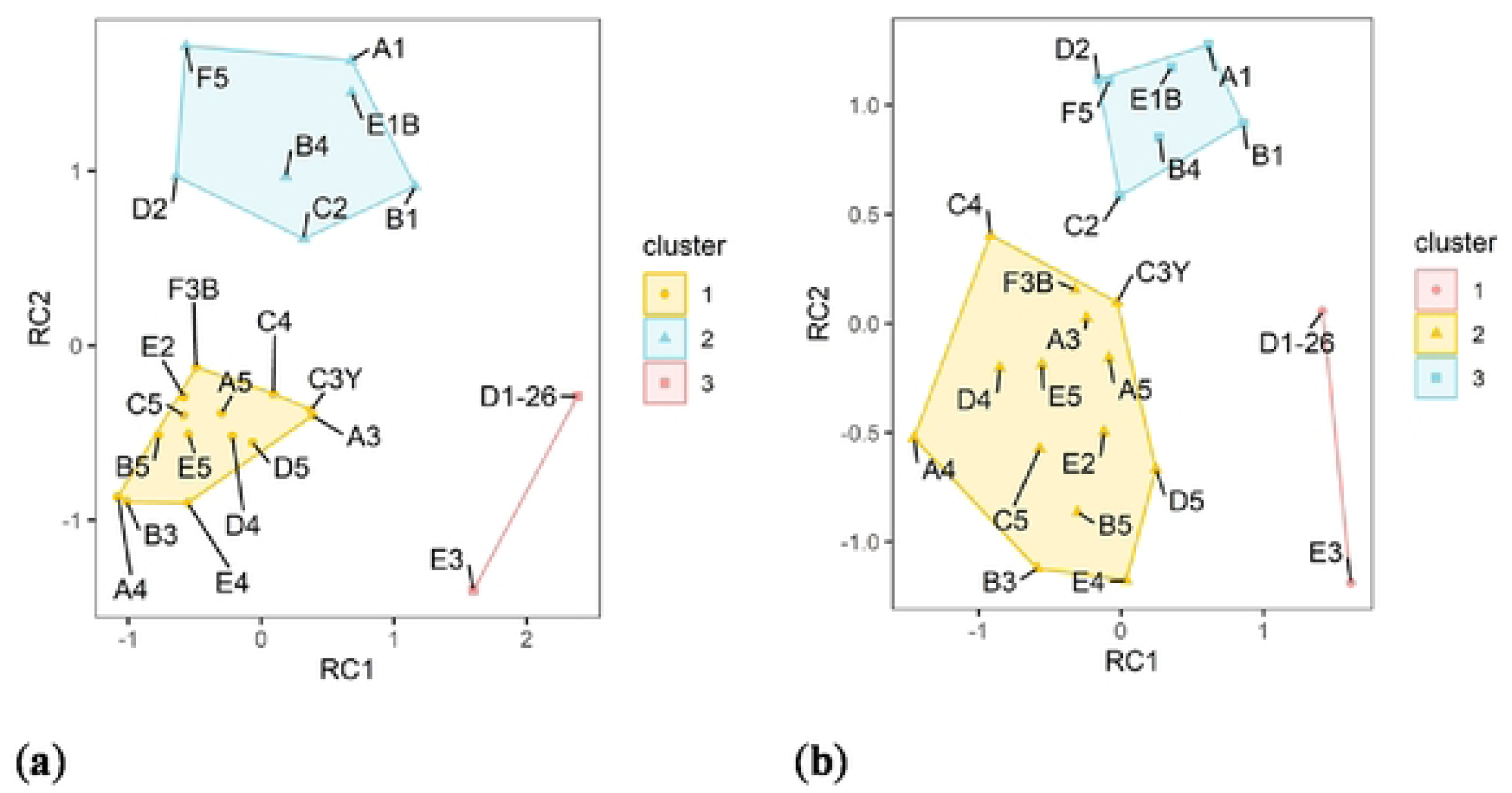
K-means clustering, k = 3: (a) applied to the results of the expert PCA to identify clusters of videos exhibiting dogs with similar emotional and motivational states; (b) applied to the results of the public PCA. Experts and public responses cluster in the same way, with each point representing one video. Videos are named after the experimental group (A-F), followed by a number and, if necessary, a letter denoting the colour of the focal dog.

**Table 5:** Sample means (x̄) and standard deviations (s) of expert scoring for each QBA term per cluster (mean scores >8 are highlighted and bold to note the leading emotional and motivational scores for videos within each cluster).

|  | Cluster 1 |  | Cluster 2 |  | Cluster 3 |  |
| --- | --- | --- | --- | --- | --- | --- |
| State | $\bar{x}$ | $s$ | $\bar{x}$ | $s$ | $\bar{x}$ | $s$ |
| Nervous/Anxious | **9** | 3.7 | 3.3 | 2.7 | 2.3 | 2.8 |
| Stressed | **8.5** | 4 | 2.7 | 2.6 | 1.9 | 2.9 |
| Relaxed | 1.3 | 1.9 | 3.7 | 3.3 | **8.8** | 4.5 |
| Comfortable | 1.4 | 2 | 5.1 | 3.9 | **8.8** | 4.2 |
| Playful | 0.7 | 1.5 | **9.7** | 4.3 | 2.1 | 2.9 |
| Interested/Curious | 4.2 | 3.7 | **9.8** | 4.2 | 5.7 | 4.4 |
| Excited | 1.4 | 2.3 | **8.9** | 3.8 | 3 | 3.6 |
| Frustrated | 3.6 | 4 | 4 | 3.3 | 0.4 | 1.4 |

While overall perception was similar, distributions of scoring among experts and members of the public differed, with members of public scoring a higher proportion of QBA terms with extreme marks of 0 or 15 (Figure 2).

**Figure 2:**
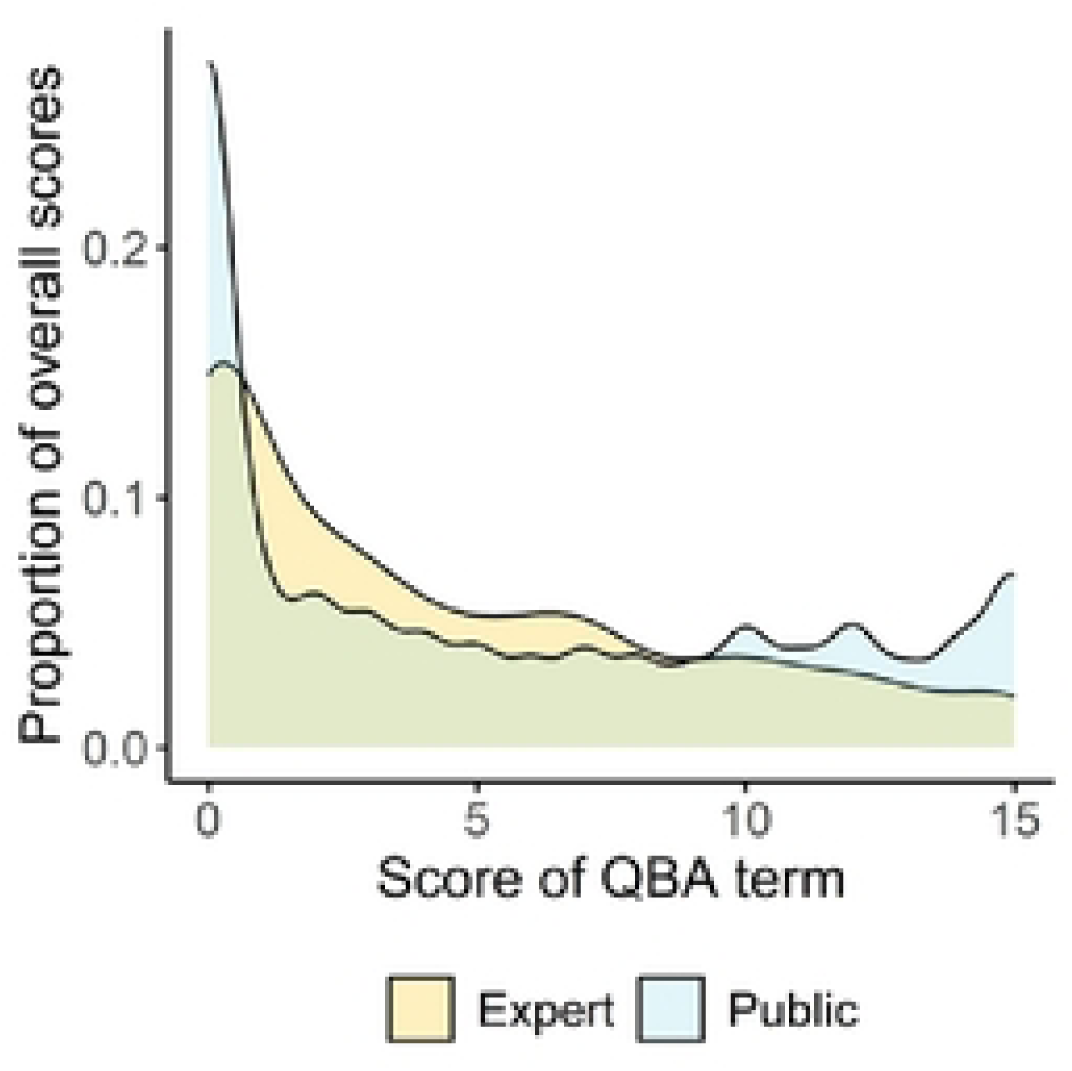
The distribution of scores (ranging from 0 to 15) by experts and the public across all videos and all QBA terms.

### 3.2 “I am uncertain” scores

A summary of IAU (“I am uncertain”) scores per QBA term and per cluster of videos for the 23 retained videos can be seen in Table. Both public (n = 15,530 video views) and expert (n = 253 video views) had the highest IAU scores for QBA terms “bored” (public, 8.7%; expert, 7.1%) and “frustrated” (public, 7.8%; expert, 5.5%). Cluster 3 (main emotional and motivational scores: “relaxed” and “comfortable”) had the largest percentage of IAU responses from experts (5%), whilst Cluster 1 (main emotional and motivational scores: “nervous/ anxious” and “stressed”) had the largest IAU percentage for public participants (4.5%). Cluster 2 (main emotional and motivational scores: “playful”, “interested/curious” and “excited”) produced the lowest IAU percentage for both groups (Table). IAU responses for excluded videos are detailed in Section II of the Supplementary Materials.

**Table 6:** Counts and percentages of “I am uncertain” (IAU) responses for each QBA term and each cluster in both expert and public survey responses. Percentages are out of 253 and 15,530 total responses for expert and public respectively for QBA terms. Sample size of QBA term responses for clusters varied due to the differing number of videos per cluster (Cluster 1 contained 14 videos, Cluster 2 seven, and Cluster 3 two).

| QBA Term | Expert IAU |  | Public IAU |  |
| --- | --- | --- | --- | --- |
|  | Count | Percentage | Count | Percentage |
| Bored | 18 | 7.1 | 1,351 | 8.7 |
| Frustrated | 14 | 5.5 | 1,213 | 7.8 |
| Comfortable | 9 | 3.6 | 496 | 3.2 |
| Excited | 8 | 3.2 | 467 | 3.0 |
| Nervous/Anxious | 6 | 2.4 | 351 | 2.3 |
| Playful | 5 | 2.0 | 525 | 3.4 |
| Interested/Curious | 5 | 2.0 | 347 | 2.2 |
| Stressed | 4 | 1.6 | 420 | 2.7 |
| Relaxed | 3 | 1.2 | 493 | 3.2 |
| Cluster 1 | 45 | 3.2 | 3,833 | 4.5 |
| Cluster 2 | 17 | 2.5 | 1,354 | 3.3 |
| Cluster 3 | 10 | 5.1 | 476 | 3.4 |

### 3.3 Scoring difficulty descriptive statistics

Figure 3 compares the experts’ and public’s subjective difficulty ratings of the three video clusters (also see Figure S1 and Section III in Supplementary Materials). Both experts and public, on average, rated videos in Cluster 2 as the easiest to score. In contrast, experts rated videos in Cluster 3, and the public rated videos in Cluster 1, as most difficult to score. Further comparisons between perceptions of difficulty in the two populations were not made, given the differences in sample size and the experts’ purpose as a reference category rather than a study population.

**Figure 3:**
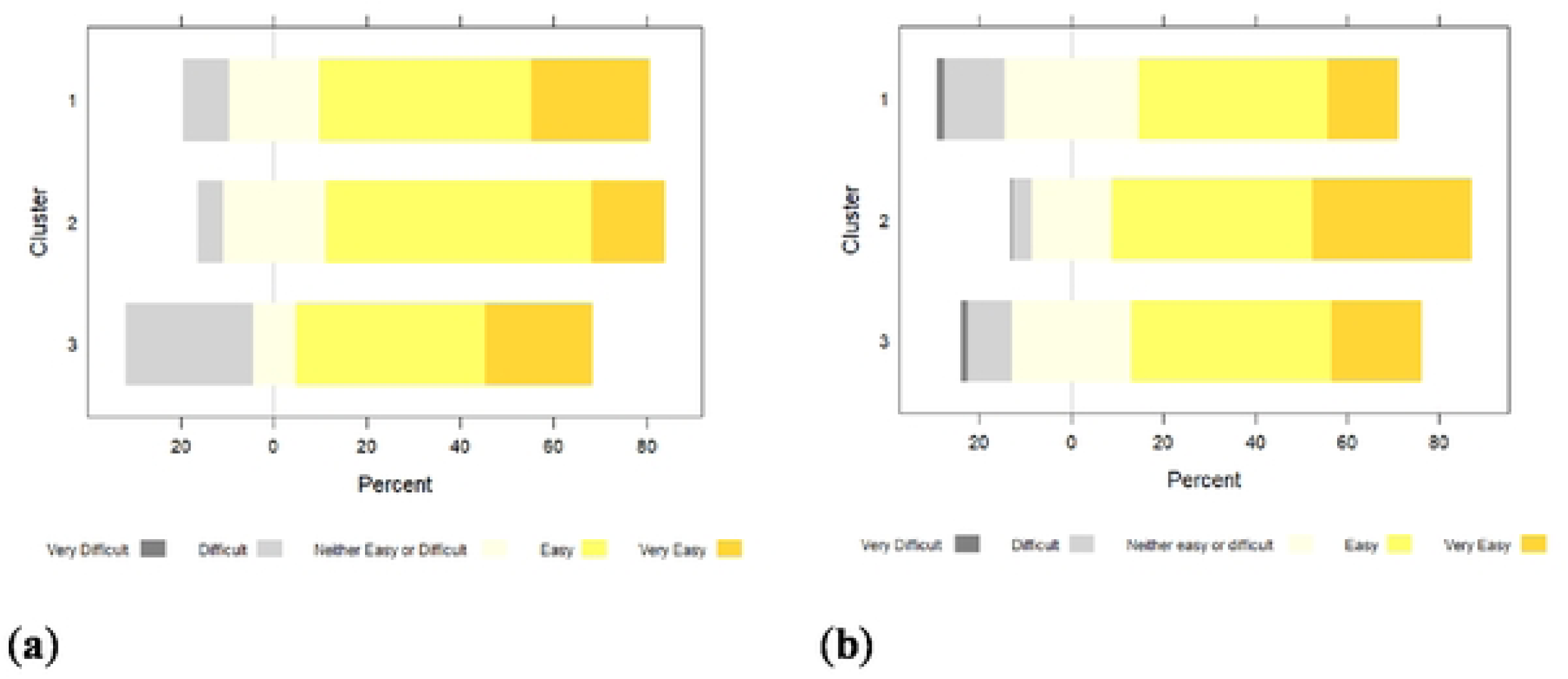
Difficulty ratings for scoring videos based on a 5-point Likert scale: (a) Experts’ difficulty ratings per cluster; (b) Public’s difficulty ratings per cluster. See Figure S1 for a by-video breakdown.

### 3.4 Emotional and motivational states associated with larger difference variable values and greater perceived difficulty in scoring

#### 3.4.1 Difficulty ratings

Videos in Cluster 2 (“playful”, “excited”, and “interested/curious” emotional/motivational states) had a lower probability (odds ratio = 0.139, CI: 0.06 – 0.33, probability estimate = 1.90%) of being classified as difficult compared to videos in Cluster 1 (“stressed” and “nervous/anxious” emotional/motivational states; odds = 0.140, CI: 0.09 – 0.23, probability estimate = 12.25%).

This means that members of the public found videos in Cluster 2 easier to score than in Cluster 1 (Model 1). The same pattern was found as RC2 increased, i.e. as the dogs were perceived as more excited and playful. A video which had a 1-point higher RC2 score (ranges from −1.4 to 1.72) had on average 0.33 times the odds of being classed as difficult compared to a video with a lower RC2 score (Model 2). See Table 2 and Table 3 for a reminder of model hypotheses and specifications, and see Supplementary Materials Section IV for more detailed results and model comparisons.

#### 3.4.2 Accuracy ratings (log of difference variable)

When participants scored all QBA terms for a video and used no “I am uncertain” scores, neither clusters nor rotated components were significantly associated with differences in scoring accuracy as measured by the log of the difference variable (Models 3 and 4, see Supplementary Materials Section IV for further details).

#### 3.4.3 Difference variable missingness models

Models 3, 4, 8, and 10 excluded observations with a missing DV from analysis. These excluded observations represented approximately 12.6% of the total sample (1,959/15,530 total observations excluded). The missingness of DV (caused by “I am uncertain” scores for QBA terms) was not random across clusters or rotated components. Videos in Cluster 1 were more likely to have missing DV values than videos in Cluster 3 based on post-hoc Tukey test, but no other significant differences were found between clusters (Model 5; Table). When assessing the relationship between the rotated components and missingness of DV (Model 6), the ratio of odds ratios for the interaction term between the rotated components (ORR = 1.218, z-value = 4.25, p < 0.001) indicates that where both RC1 and RC2 were at either the far end of the upper (positive) or lower (negative) range, the likelihood for missing DV values increased. In the opposite quadrants (negative RC1 and positive RC2 or the reverse), we see the reverse, with a lower likelihood of missing DV values. These quadrants are associated with dogs who are playful/excited (negative RC1, positive RC2) and relaxed/comfortable (positive RC1, negative RC2, Model 6). This means that participants were more likely to have scored “I am uncertain” for a QBA term when dogs exhibited RC1 and RC2 scores that were both highly positive (few dogs fell within this range) or both negative (stressed/anxious dogs). Please see Supplementary Materials Section IV for detailed results of missingness analyses.

**Table 7:** Model 5 post-hoc comparison of DV missingness by Cluster with Tukey-adjusted p-values. Details of Model 5 results can be found in Section IV of Supplementary Materials.

| Contrast | Odds Ratio | SE | Z-ratio | p-value |
| --- | --- | --- | --- | --- |
| Cluster 1 / Cluster 2 | 1.14 | 0.075 | 1.983 | 0.12 |
| Cluster 1 / Cluster 3 | 1.36 | 0.144 | 2.929 | **0.01** |
| Cluster 2 / Cluster 3 | 1.20 | 0.138 | 1.557 | 0.26 |

### 3.5 Expert and public subjective assessment of difficulty and competence in scoring

#### 3.5.1 Average expert difficulty rating vs. expert agreement by video (ICC estimate)

A spearman’s rank order correlation indicates that there was a negative monotonic relationship between the average expert difficulty rating and the video’s intraclass correlation coefficient estimate (Analysis 7: ⍴(24) = −0.452, p < 0.05). As experts on average rated videos to be more difficult, the level of agreement between experts on what that video was showing also decreased.

#### 3.5.2 Scoring Accuracy (log of public DV scores and DV missingness) as predicted by video difficulty and individual self-rated competency

Public participants’ difficulty ratings of videos were not associated with scoring accuracy when participants scored all QBA terms (Model 8). As participant-rated competence (Likert scale response to the statement “I feel confident interpreting any dog’s behaviour”) increased, the likelihood of the DV score being missing decreased (OR = 0.84, z-value = −3.42, p < 0.001) and as participant-rated difficulty increased, the likelihood of the DV score being missing increased (OR = 1.48, z-value = 10.64, p < 0.001). These findings indicate that participants who rated themselves as more competent were less likely to have missing DV scores as they were less likely to score QBA terms as “I am uncertain” (Model 9). Where participants scored all QBA terms, the average of the experts’ difficulty ratings for each video was correlated with public scoring accuracy (Model 10). Specifically, a model incorporating fixed effects of average expert difficulty rating, member of the public’s self-rated competence, and an interaction of expert difficulty rating and self-rated competence, along with random effects of video ID and participant ID outperformed all simpler models (Table S6, Supplementary Materials). The model’s marginal R-squared value indicates a higher proportion of the variance is explained by the fixed effects of Model 10 than by those of Model 8. However, Model 10 still only accounts for approximately 29% of the total variance in log DV score, with the fixed effects of individual public competency, average expert difficulty rating, and their interaction still only accounting for approximately 10% of the total variance (Table S7, Supplementary Materials). Based on Model 10, there may also be a significant interaction between the average of experts’ difficulty ratings and public participants’ self-rated confidence at interpreting any dog’s behaviour (Table 8), with more confident participants performing worse the more difficult videos became according to the experts. See Supplementary Materials Section IV for details.

**Table 8:**
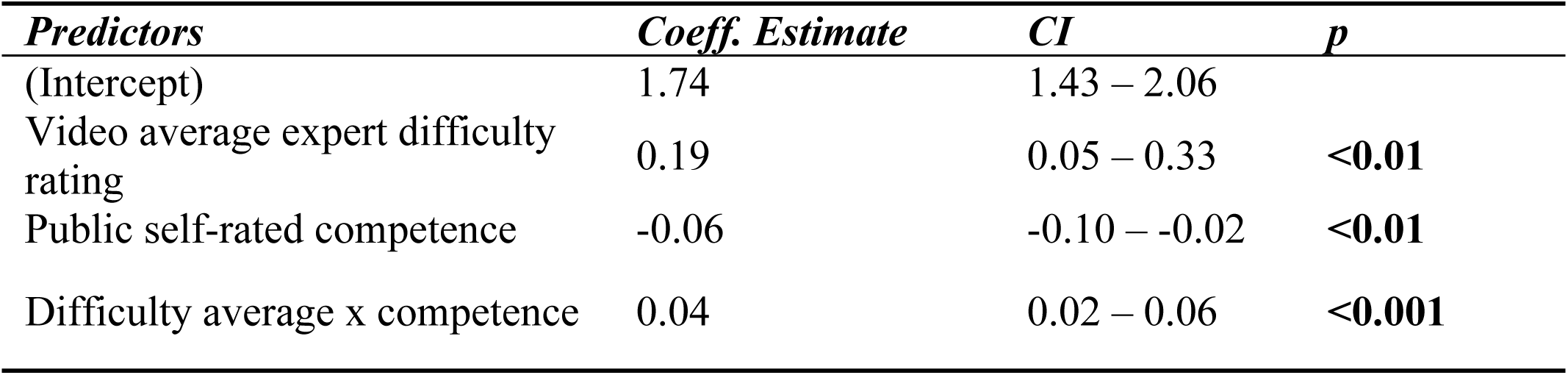
Coefficient estimates, confidence intervals, and p-values for all fixed-effect predictors included in Model 10.

Predicted DV increased (indicating worse performance in public scoring) as the average of the expert’s difficulty rating of videos increased (Figure S2, Supplementary Materials). There was a slight effect of public participant self-rated competence, with an interaction with expert difficulty rating. Public participants who considered themselves very competent scored better on “very easy” videos. However, those participants scored worse than participants who rated themselves as less competent as video difficulty increased. The lack of a significant effect of individual public difficulty rating in Model 10 may have also been due to an inability to calculate DV for observations where a member of the public did not rate all QBA terms and instead selected “I am uncertain”.

## 4 Discussion

Our findings centre on three main themes: first, the comparison of the overall perception of dog emotional and motivational states in videos between experts and the public; second, the identification of emotional and motivational states that are perceived as more and less difficult to interpret by experts and the public; and third, the assessment of the accuracy with which the public and experts evaluate their own competence and the difficulty in interpreting emotional and motivational states of dogs from silent videos.

### 4.1 Overall perception

Public participants demonstrated a general ability to interpret dog behaviours when compared to evaluations carried out by an expert panel. Similar PCA loadings for public and expert results on the two PCA components and identical video clustering for each group support this conclusion. These findings complement those of Bloom and Friedmann (2013) who found that both experts (dog trainers) and laypeople could correctly assign a predominant emotion to a dog’s facial display from a photo. The strong similarity in loadings for both the public and expert PCAs suggests a consistent interpretation of the QBA terms by both groups. However, plotting of score distributions for expert and public participants showed that the public tended to assign extreme scores (0 and 15) proportionally more frequently. Members of the public might be less adept at identifying emotional states where valence and arousal are not at extremes: possibly, missing subtle behaviour signals. Past research has shown differences in brain activation during observations of social behaviour of dogs between experts and non-experts (Kujala et al., 2012), suggesting training-mediated changes in perceptual focus among experts.

PCA provided insights into the overall perception of dog emotional and motivational states, with one rotated component primarily associated with emotional valence and the second with arousal level. The first rotated component loaded heavily on the fear-comfort continuum and included “nervous/anxious”, “stressed”, “relaxed”, and “comfortable” terms; the second component loaded for “playful”, “interested/curious”, “excited”, and on a lesser level “nervous/anxious” and “stressed” terms. The identified pattern is similar to the dimensional models of emotions (e.g., (Barrett, 2006; Barrett et al., 2007; Carver, 2001; Posner et al., 2005; Watson et al., 1999)), where the emphasis is on emotional dimensions that emerge from a combination of valence of emotions (positive or negative) and arousal levels (Mendl et al., 2010), rather than focusing on basic emotions as the fundamental components of emotional experience.

### 4.2 Emotional states – Identification and ease of interpretation

Public participants perceived the emotional and motivational states of Cluster 2, i.e., playful, excited, and curious dogs, as easier to interpret than those of Cluster 1, i.e., fearful, stressed, and/or anxious dogs (Models 1 and 2; IAU score comparisons), aligning with previous research (Kerswell et al., 2009, 2013; Lakestani et al., 2014; Tami & Gallagher, 2009; Wan et al., 2012). Members of the public were generally less certain when interpreting dogs exhibiting more negatively valenced emotional and motivational states (IAU scores and Models 1, 2, 5, and 6). However, this difference was not reflected in actual scoring ability when participants were willing to score all QBA terms, with no difference in scoring accuracy between videos of nervous and anxious dogs versus playful, curious, and excited dogs (Models 3 and 4).

Public participants demonstrated a heightened awareness of their knowledge gaps concerning negative emotional and motivational states (higher perceived difficulty ratings and more frequent IAU scores), yet they exhibited potential overconfidence regarding their ability to interpret dogs exhibiting positively valenced emotional states. This observation is consistent with Kerswell et al.’s reports that puppy owners rated their confidence as higher in interpreting more positively valenced emotional states, but that the number of behavioural cues they could enumerate for these emotional states did not surpass the number they could recall for negatively valenced emotional states. Importantly, negative emotional and motivational states are more likely to be associated with concerning behaviours, such as aggression. Consequently, limited ability to recognise when a dog is experiencing negative emotional and motivational states can also increase the risk of a negative human-dog interaction. This may manifest in scenarios where individuals are unable to recognise a dog requires space, reassurance, or tailored handling approaches.

### 4.3 Boredom and frustration prove difficult

Low inter-expert reliability, high proportions of “I am uncertain” scores, and exclusion from the expert PCA due to low sampling adequacy according to KMO tests support the conclusion that “boredom” and “frustration” were terms that participants (both expert and public) found difficult to interpret and score. This may be because frustration – defined as an animal’s reaction after surprising incentive omissions (Papini & Dudley, 1997) – could occur in scenarios featuring both positively and/or negatively valenced emotional and motivational states. Lack of sound and contextual information may have made frustration particularly difficult to identify, as vocalizations are frequently cited as behavioural cues of frustration in dogs, and humans have been shown to pay attention to vocalizations when differentiating between canine emotional and motivational states (Jakovcevic et al., 2013; Lakestani et al., 2014; Pongrácz et al., 2005, 2006; Volodina et al., 2006). Chronic boredom, sometimes demonstrated as “waking inactivity,” can lead to distress and poor welfare and is associated with behavioural problems (Burn, 2017; Harvey et al., 2019). Behavioural similarities between boredom and depression have been noted (Harvey et al., 2019), and frustration often leads to signs of distress similar to those elicited by fear (Jakovcevic et al., 2013), emphasising the relevance to dog welfare of identifying both states.

### 4.4 Assessing difficulty and competence

Experts and members of the public assessed video difficulty and their own competence in assessments differently. The average expert difficulty rating for videos was correlated both with the level of inter-expert agreement (Analysis 7) and with how well the public scored the QBA terms (Model 10), indicating experts were adept at estimating video difficulty. In the public dataset, a higher difficulty rating by a participant was associated with an increased likelihood of “I am uncertain” scores (Model 9), but where participants scored all QBA terms, there was no relationship between the public’s difficulty perception and scoring accuracy (Model 8).

Similarly, while the public rated videos in Cluster 2 (playful, excited, and curious dogs) as easier to score than those in Cluster 1 (anxious, stressed, and fearful dogs) and videos with higher RC2 scores as easier than those with lower RC2 scores, these differences in perceived difficulty were not reflected in scoring accuracy when participants used no “I am uncertain” scores (Models 1 – 4).

Members of the public assessed their own competence poorly, measured as their reported general self-confidence in interpreting any dog’s behaviour (Models 8 and 10). While public participants with high overall self-confidence scored well on easy (as rated by experts) videos, their performance on more difficult videos tended to be worse than that of less confident participants. Past research showed that confidence in dog behaviour interpretation is shaped by a person’s experience with dogs (Kerswell et al., 2009, 2013; Wan et al., 2012). Paradoxically, more experience has in some studies been associated with poorer interpretation of dog behaviour, possibly due to experience biasing the assessment towards positive interpretations of dog behaviour (Bloom & Friedman, 2013) or due to overconfidence. This potential disconnect between self-perceived competence and actual ability should be considered when developing human behaviour change strategies. Interventions aimed at improving human-dog interactions should assume that competence, confidence, and ability to evaluate one’s competence, regarding interpretation of dog emotions, are all factors that need to be addressed to facilitate change.

### 4.5 Limitations

Videos were provided without sound to avoid acoustic cues, such as comments from observers, biasing participants’ perception. However, other environmental and holistic cues may have led to some videos being “easier” to interpret or more vulnerable to bias (for example, the presence of toy interaction). Whilst a large effort was made to capture a representative public participant sample, a common survey recruitment trend occurred: participant bias towards a middle aged, female audience (also noted in other studies e.g., (Meints & de Keuster, 2009; Westgarth & Watkins, 2015)). Male participants were evenly spread across all video groups and all video clusters. Past research (Mariti et al., 2012) has shown gender differences in perception of pet stress, emphasising the importance of continued attempts to address this imbalance in future research on public perception of dog behaviour and emotion.

A participant’s personality (Kujala et al., 2017), cultural milieu (Amici et al., 2019), and beliefs about traits or other characteristics (Menchetti et al., 2018), along with a dog’s breed (Gunter et al., 2016), size (Arhant et al., 2010), framing of information during an experiment (Wells et al., 2012), and the physical environment (Wells et al., 2012) can influence behavioural interpretations. Cultural milieu and contextual information remained mostly constant across participants in this study as all participants resided in the UK/ROI. Our study included a range of dog breeds and sizes, randomly distributed across video groups. This survey did not explore empathy and other personality traits that may impact an individual’s perception and response to animal behaviours and emotion. Finally, the study did not compare the expert-based assessment with animal-based measures of welfare. Further work is required to validate expert-based assessments against animal-based measures for a range of emotional and motivational states.

## 5 Conclusions

This study identified strong similarities in the overall perception of dog emotional and motivational states between the public and experts. Public and experts’ uncertainty in identifying behavioural signs indicative of boredom and frustration highlights a need for further research and education, particularly as both states are highly relevant to dog welfare. Public participants perceived videos of dogs exhibiting stressed, anxious, and fearful behaviour as more difficult to interpret and were less likely to score all QBA terms for these videos. The discrepancy identified between the public’s perception of their own competence, the perceived difficulty of interpretation, and their scoring accuracy emphasizes the importance of addressing both knowledge gaps and instances of overconfidence to improve human-dog interactions.

## Author Contributions (authors in alphabetic order by category)

Conceptualization, N.H., J.M., and L.S.; methodology, N.H., J.M., J.M.M., S.O.G, and L.S.; validation, J.M.M., and L.S.; formal analysis, J.M.M., L.S.; investigation K.G., N.H., and L.S.; resources, N.H., J.M., S.O.G., and M.U.; data curation, J.M.M. and L.S.; writing—original draft preparation, N.H., J.M.M. and L.S.; writing—review and editing, J.M., J.M.M., S.O.G., L.S., and M.U.; visualization, J.M.M. and L.S.; supervision, N.H. and S.O.G.; project administration, L.S.; funding acquisition, N.H., J.M., and M.U. All authors have read and agreed to the published version of the manuscript.

## Institutional Review Board Statement

The study was conducted in accordance with the Declaration of Helsinki and approved by the Dogs Trust Ethical Review Board (reference number ERB41, approved on 21 January 2021).

## Informed Consent Statement

Informed written consent was obtained from all study participants and from all owners or rehoming centre staff who recorded videos to be used in the study.

## Acknowledgments

We would like to thank members of Dogs Trust’s community who contributed or allowed us access and permissions to videos for use in the study, the eleven Dogs Trust behaviour experts who completed the expert version of the study survey, and Dogs Trust Communications and Digital Directorate who guided and assisted us with survey distribution and advertisement. Thank you, Kirsten McMillan, for your helpful manuscript comments. Finally, we thank Charlotte Huggins and Tamsin Durston, both experts in canine behaviour, who assisted with the creation of the video tutorials shown to participants after survey completion.

